# Improving Nitrofurantoin Resistance Prediction in *Escherichia coli* from Whole Genome Sequence by Integrating NfsA/B Enzyme Assays

**DOI:** 10.1101/2024.01.25.577238

**Authors:** Punyawee Dulyayangkul, Jordan E Sealey, Winnie WY Lee, Naphat Satapoomin, Carlos Reding, Kate J. Heesom, Philip B Williams, Matthew B Avison

## Abstract

Nitrofurantoin resistance in *Escherichia coli* is primarily caused by mutations damaging two enzymes, NfsA and NfsB. Studies based on small isolate collections with defined nitrofurantoin MICs have found significant random genetic drift in *nfsA* and *nfsB* making it extremely difficult to predict nitrofurantoin resistance from whole genome sequence (WGS) where both genes are not obviously disrupted by nonsense or frameshift mutations or insertional inactivation. Here we report a WGS survey of 200 *E. coli* from community urine samples, of which 34 were nitrofurantoin resistant. We characterised individual non-synonymous mutations seen in *nfsA* and *nfsB* among this collection using complementation cloning and assays of NfsA/B enzyme activity in cell extracts. We definitively identified R203C, H11Y, W212R, A112E, A112T and A122T in NfsA and R121C, Q142H, F84S, P163H, W46R, K57E and V191G in NfsB as amino acid substitutions that reduce enzyme activity sufficiently to cause resistance. In contrast, E58D, I117T, K141E, L157F, A172S, G187D and A188V in NfsA and G66D, M75I, V93A and A174E in NfsB, are functionally silent in this context. We identified that 9/166 (5.4%) of nitrofurantoin susceptible isolates were “pre-resistant”, defined as having loss of function mutations in *nfsA* or *nfsB*. Finally, using NfsA/B enzyme activity assay and proteomics we demonstrated that 9/34 (26.5%) of nitrofurantoin resistant isolates carried functionally wild-type *nfsB* or *nfsB*/*nfsA*. In these cases, enzyme activity was reduced through downregulated gene expression. Our biological understanding of nitrofurantoin resistance is greatly improved by this analysis, but is still insufficient to allow its reliable prediction from WGS data.

## Introduction

Nitrofurantoin is the first choice for treatment of uncomplicated urinary tract infections in adults in the United Kingdom, of which a majority are caused by *Escherichia coli* (1). Nitrofurantoin enters *E. coli* and the enzymes NfsA or NfsB catalyse its degradation to release toxic free radicals that kill. Nitrofurantoin resistance is relatively uncommon compared with other potential agents (e.g. trimethoprim[1]), and primarily involves mutations disrupting the functions of both NfsA and NfsB (2–6).

In a recent study of 9 nitrofurantoin resistant urinary *E. coli* from London, United Kingdom, whole genome sequencing revealed 9 different sequence types, and a variety of *nfsA* and *nfsB* mutations including nonsense mutations, frameshifts and some missense mutations, plus insertional inactivation with various IS elements (6). Gene loss has also been identified in laboratory selected mutants (7). However, wider analysis of whole genome sequence data for *E. coli* clinical isolates has revealed that *nfsA* and *nfsB* are very diverse, with significant random genetic drift, and it is difficult to accurately predict nitrofurantoin resistance unless both genes are clearly disrupted (2–6).

One hypothesis to explain the low prevalence of nitrofurantoin resistance in *E. coli* in the UK despite widespread nitrofurantoin use is that individual *nfsA* or *nfsB* mutations do not give a sufficiently strong selective advantage when nitrofurantoin is dosed properly (8). However, resistant isolates have emerged (2–6), and there is some evidence of nitrofurantoin susceptible isolates from human and other sources having a loss of function mutation in one of the two genes (6). Collections of sequenced bloodstream isolates are numerous (6), but without an accurate way of predicting which amino acid changes seen in NfsA or NfsB are due to random genetic drift, and which damage the functions of these two enzymes, it remains extremely difficult to accurately predict nitrofurantoin susceptibility/resistance status of such isolates (6). Since nitrofurantoin is seldom tested against bloodstream *E. coli* in the laboratory, because it has no utility for the treatment of systemic infections (1), phenotypic data are not readily available (6). Therefore, the ability to predict resistance from whole genome sequence data would be extremely useful. For example, this would allow us to investigate whether nitrofurantoin “pre-resistance”, defined by the loss of NfsA or NfsB activity but where phenotypic resistance has not been achieved because the other enzyme remains functional, is associated with an increased incidence of treatment failure for urinary tract infection, and an increased incidence of pyelonephritis and urosepsis.

The aim of the work reported here was to establish methods that can be used to address these clear gaps in our understanding of how NfsA/B activity can be predicted from whole genome sequence data. We established a complementation cloning methodology to differentiate genetic drift from loss of function mutations. The mutations studied were those identified from genomic surveillance of urinary *E. coli* (with confirmed nitrofurantoin resistance/susceptibility status) submitted from primary care to a diagnostic laboratory serving a population of approximately 1.5 million people. We have also used a simple enzymatic assay that can measure NfsA and NfsB activity in bacterial cell extracts. This assay, coupled with the use of LC-MS/MS proteomics analysis revealed that NfsA and/or NfsB production can be downregulated without deviation from wild-type NfsA/B enzyme function, a phenomenon that we found to be common. This discovery adds additional complexity to the implementation of any proposed use of whole genome sequence analysis to predict nitrofurantoin susceptibility/resistance. We suggest that rapid in vitro NfsA/B enzyme assays might have more utility to do this until our biological understanding of genotype to nitrofurantoin resistance phenotype relationships is more complete.

## Results

### In-patient isolate clonality but no evidence of person to person transmission of nitrofurantoin resistant urinary E. coli from primary care

In 2020, the prevalence of nitrofurantoin resistance among isolates from 17,581 *E. coli*-positive urine samples submitted from primary care to the Severn Pathology diagnostic laboratory, North Bristol NHS Trust, was 1.34%. During September and October 2020, nitrofurantoin resistant *E. coli* were collected. After confirming resistance using broth microdilution and removing all but the first isolate from each patient during the surveillance period, 34 isolates were sent for whole genome sequencing. Twenty were susceptible to the other 5 antibacterials tested during routine diagnostics. Of the remainder, 8 were cefalexin resistant, of which 7 cefpodoxime resistant; 8 were ciprofloxacin resistant; 5 were amoxicillin/clavulanate resistant; 10 were gentamicin resistant (**Table 1**).

**Table 1:** Characterisation of 34 deduplicated nitrofurantoin resistant urinary *E. coli* from primary care. LEX, cefalexin; CPD, cefpodoxime; CIP, ciprofloxacin; AMC, amoxicillin/clavulanate; GEN, gentamicin; NIT, nitrofurantoin; S, sensitive; I, intermediate; R, resistant; green, wild-type like function; orange, function not individually tested; red, loss of function.

| Isolate Code | Susceptibility to |  |  |  |  |  | Sequence Type | Effect of non-synonymous mutations relative to *E. coli* K-12 |  |
| --- | --- | --- | --- | --- | --- | --- | --- | --- | --- |
|  | LEX | CPD | CIP | AMC | GEN | NIT |  | NfsA | NfsB |
| 39675 | S | S | S | S | S | R | ST14 | I117T, K141E, G187D, W212R | Del203-207 |
| 39677 | S | S | S | S | S | R | ST404 | 187FS | G66D, V93A, A174E |
| 39679 | S | S | S | S | S | R | ST404 | 113Stop | 101Stop |
| 39680 | S | S | S | S | S | R | ST23 | deletion between *rimK* and *rcdA* | 133Stop |
| 39681 | S | S | S | S | S | R | ST95 | 17FS | Insertionally inactivated after K14 |
| 39683 | S | S | S | S | S | R | ST73 | 193Stop | 48FS |
| 39684 | S | S | S | S | S | R | ST95 | 20Stop | G66D, V93A, Q142H, A174E |
| 39686 | S | S | S | S | S | R | ST10222 | I117T, K141E, G187D, R203C | Insertionally inactivated after K14 |
| 39688 | S | S | S | S | S | R | ST131 | 67Stop | G66D, V93A, P163H, A174E |
| 39969 | R | R | R | I | R | R | ST131 | 92FS | G66D, F84S, V93A, A174E |
| 39985 | R | R | R | R | S | R | ST405 | 32FS | 24FS |
| 39995 | R | R | R | R | S | R | ST443 | L157F | M75I, V93A |
| 40001 | R | R | R | S | R | R | ST131 | H11Y, I117T, K141E, G187D | W46R, G66D, V93A, A174E |
| 40002 | R | R | R | R | R | R | ST131 | 68FS | 181Stop |
| 40103 | S | S | S | S | S | R | ST73 | I117T, K141E, G187D, A188V, R203C | deletion between *ybdF* and *pheP* |
| 40104 | S | S | S | S | S | R | ST404 | A112E, I117T, K141E, G187D | G66D, V93A, A174E |
| 40105 | S | S | S | S | R | R | ST131 | 58Stop | K57E, G66D, V93A, A174E |
| 40107 | S | S | S | S | R | R | ST131 | A34V, A112T, I117T, K141E, G187D | G66D, V93A, A174E |
| 40108 | S | S | S | S | S | R | ST58 | 212Stop | Del95-105 |
| 40109 | S | S | S | S | S | R | ST69 | 100Stop | V93A, R121C |
| 40110 | S | S | S | S | S | R | ST69 | E58D, I117T, K141E, A172S, G187D, R203C | Insertionally inactivated after K14 |
| 40111 | S | S | S | S | S | R | ST998 | Ins129I | D66G, E137Q, E165K, L172F, A174E |
| 40113 | S | S | S | S | S | R | ST5281 | deletion between *ybjN* and *aspT* | 36Stop |
| 40114 | S | S | S | S | S | R | ST58 | deletion between *nfsA* and *ybjO* | 110FS |
| 40115 | S | S | S | S | R | R | ST131 | deletion between *nfsA* and *gsiC* | G66D, V93A, A174E |
| 40116 | S | S | S | S | S | R | ST73 | 204Stop | G66D, V93A, A174E |
| 40118 | S | S | S | S | S | R | ST95 | 214FS | deletion between *ybdF* and *mscM* |
| 40119 | S | S | S | S | S | R | ST14 | Ins189LAQYDEQ | G66D, V93A, A174E, V191G |
| 40121 | S | S | S | S | R | R | Novel ST | A34V, A112T, I117T, K141E, G187D | deletion between *ybdF* and *pheP* |
| 53852 | S | S | R | S | R | R | ST131 | 72Stop | deletion between *leuC* and *ybcH* |
| 53866 | R | S | R | R | S | R | ST744 | WT | WT |
| 53898 | S | S | R | S | R | R | ST1193 | I117T, K141E, E178G, G187D | G66D, V93A, A174E |
| 55968 | R | R | S | S | S | R | ST450 | 55FS | V93A |
| 55971 | R | R | S | R | R | R | ST73 | 141Stop | 101Stop |

WGS revealed remarkable diversity among the 34 deduplicated nitrofurantoin resistant urinary isolates, both phylogenetically and in terms of the mobile resistance genes and NfsA/B amino acid variations identified (**Figure 1**, **Table 1**). There were no instances of the same isolate (or even one with the same combination of NfsA/B variants) being isolated from urine samples provided by two people. However, sequencing of additional nitrofurantoin resistant isolates from four people where multiple samples were positive during our surveillance period revealed that in all cases (one person, 3 samples; three people, 2 samples each) the bacteria sequenced from each sample from the same person were essentially identical, with core genome SNP distances of <5 between them and the same acquired resistance genes and NfsA/B variations.

**Figure 1.**
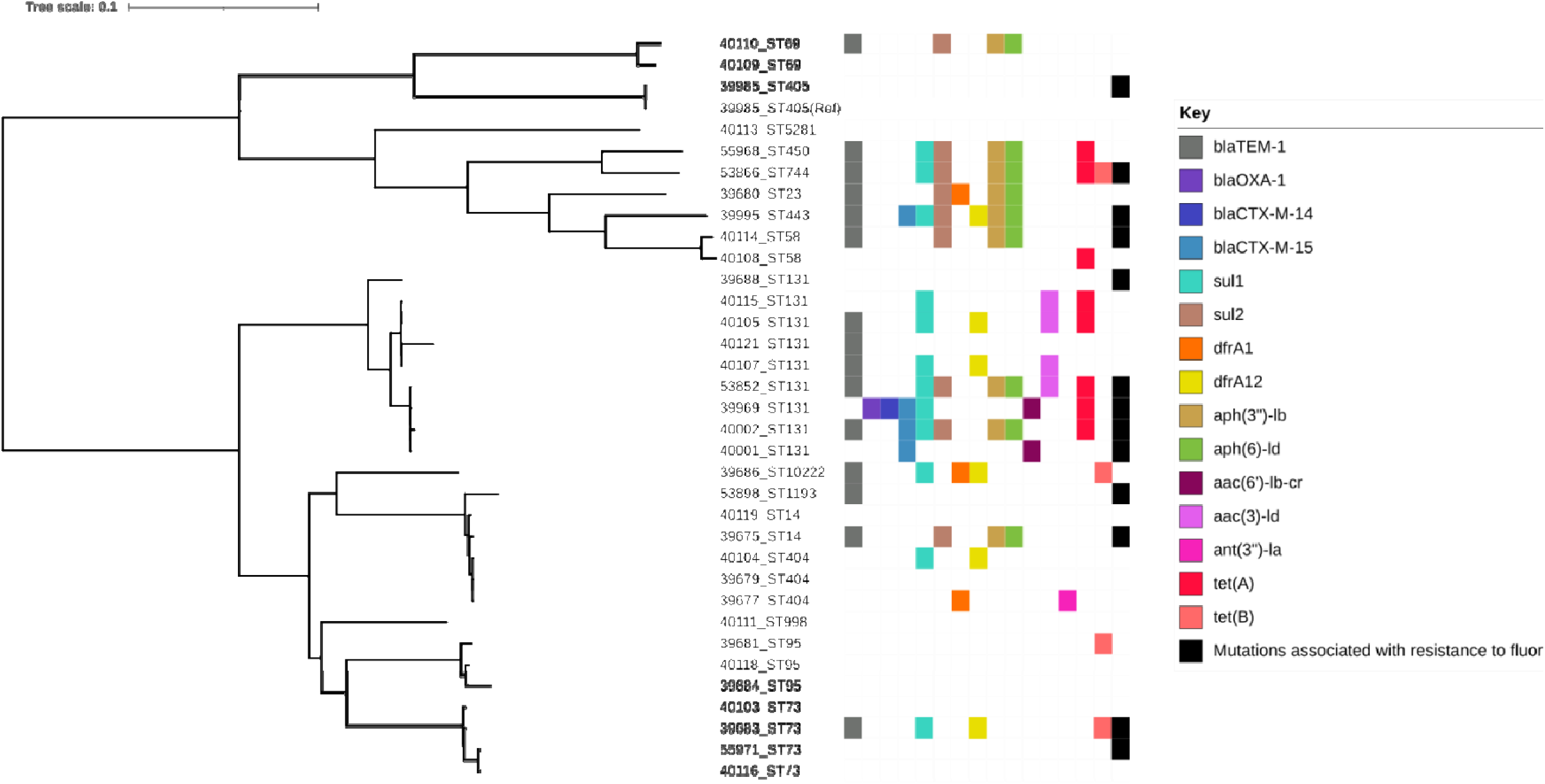
Maximum likelihood phylogenetic tree derived from a core genome alignment of whole genome sequence for 34 nitrofurantoin resistant *E. coli* from primary care urine samples. Five-digit isolate numbers are provided alongside each isolate’s sequence type (ST) and the resistance genes/mutations carried (other than those causing nitrofurantoin resistance. The reference (ref) genome is marked.

### Complementation cloning and phenotypic assays to differentiate functionally wild-type from reduced function E. coli NfsA and NfsB variants

Only 12/34 (35.2%) nitrofurantoin resistant isolates had obvious loss of function in both NfsA and NfsB (defined as nonsense mutation or frameshift in, insertional inactivation or deletion of the encoding gene), which is required for nitrofurantoin resistance without other mechanisms being involved (2–6). Additionally, 11/34 (32.4%) and 5/34 (14.7%) of nitrofurantoin resistant isolates, had obvious loss of function in only one of NfsA or NfsB, respectively (**Table 1**). Missense mutations causing loss of function were suspected in the remainder, and in many cases, missense mutations altering the amino acid sequence of NfsA or NfsB were identified, relative to *E. coli* K-12 as the wild-type reference (**Table 1**). But given prior knowledge of random genetic drift in these genes (2–6) it was considered important to account for drift using a wider collection of nitrofurantoin susceptible urinary isolates and in vitro analysis of NfsA/B variant function. Firstly, therefore, in addition to the 34 nitrofurantoin resistant isolates (**Table 1**) we sequenced 166 confirmed nitrofurantoin susceptible urinary *E. coli* isolates collected in parallel, to make a total of 200 isolates sequenced for this study (**Table S1**). It is important to note that these isolates were collected during surveillance of resistance to cefpodoxime, ciprofloxacin and amoxicillin/clavulanate in urinary *E. coli* from the community so the nitrofurantoin susceptible isolates sequenced here were resistant to at least one of these agents. Using *E. coli* K-12 as the wild-type reference, we collated amino acid sequence variant types for NfsA and NfsB in the 166 nitrofurantoin susceptible isolates (**Table 2**; **Table 3; Table S1**). Clear loss of function mutations (as defined above) were identified in 7 nitrofurantoin susceptible urinary *E. coli* isolates; 4 losing the function of NfsA (**Table 2**) and 3 losing that of NfsB (**Table 3**). No isolate had obviously lost both, as expected given that they are nitrofurantoin susceptible. Otherwise, there was clustering around 4 NfsA (including our chosen wild-type) and 6 NfsB types, with 4 additional NfsA types being represented by 1 isolate each (**Table 2**, **Table 3**). Whilst others have also reported the presence of some of these NfsA/B variant types in phenotypically nitrofurantoin susceptible isolates (2–6), in order to independently confirm variant functionality, as well as to test the functions of NfsA/B missense variants seen in the nitrofurantoin resistant isolates (**Table 1**), we set up a complementation assay, and enzymatic assays for NfsA and NfsB.

**Table 2:** Variants of NfsA seen among 166 nitrofurantoin susceptible urinary *E. coli* from primary care based on whole genome sequence.

| NfsA Variant Type<br>(susceptible isolate) | Sequence | Number of<br>isolates |
| --- | --- | --- |
| WT | none | 15 |
| 1 | I117T, K141E, G187D | 105 |
| 2 | I117T, K141E, G187D, A188V | 24 |
| 3 | E58D, I117T, K141E, A172S, G187D | 14 |
| 4 | E58D, I117T, K141E, G187D | 1 |
| 5 | G187D | 1 |
| 6 (Loss of Function) | A112E (missense), I117T, K141E, G187D | 1 |
| 7 (Loss of Function) | K141E, G187D, T202P (missense) | 1 |
| Loss of Function | Group 1 + K141* (nonsense) | 1 |
| Loss of Function | 228FS (frameshift) | 1 |
| Loss of Function | Deletion *rimK* to *rcdA* | 1 |
| Loss of Function | Insertional inactivation with IS26 | 1 |

**Table 3:**
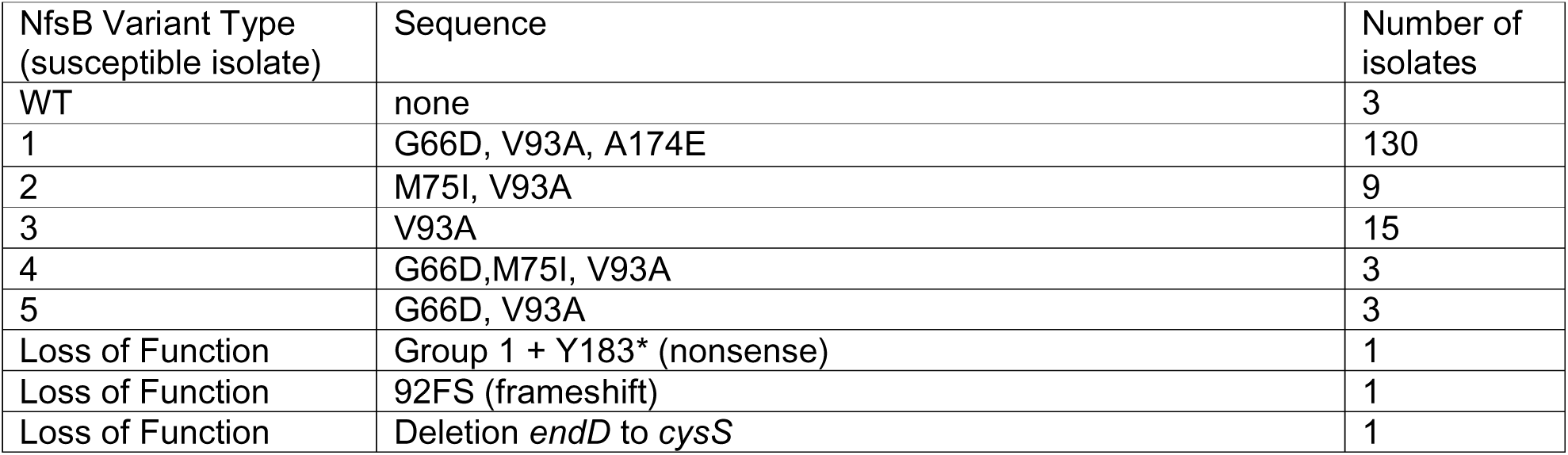
Variants of NfsB seen among 166 nitrofurantoin susceptible urinary *E. coli* from primary care based on whole genome sequence.

Using nitrofurantoin resistant urinary isolate 39681, which has a frameshift in *nfsA* and insertional inactivation of *nfsB* (**Table 1**) as a recipient for complementation, we introduced plasmid pK18 with *nfsA* or *nfsB* variants inserted, in each case with its own ribosome binding site. Expression of the inserted gene was from the leaky *lac* promoter on the vector. Since loss of function in both NfsA and NfsB is required to confer nitrofurantoin resistance (2–6), *nfsA* or *nfsB* alleles that encode functional enzymes will complement isolate 39681 to nitrofurantoin susceptibility. Non-functional variants will not. To give a more accurate measure of enzyme function, we also devised enzyme assays for NfsA and NfsB by monitoring nitrofurantoin breakdown in the presence of cofactors NADH (specific for NfsB [9-11]) or NADPH (used by both NfsA and NfsB [9-11]) upon addition of crude bacterial cell extract from 39681 recipient carrying the various recombinant *nfsA*/*B* plasmids to the reaction.

Using this approach, we were able to screen all the variants found among our 200 sequenced urinary isolates (**Tables 1 and S1**). This enabled us to confirm that the following NfsA amino acid substitutions do not alter function sufficiently to confer nitrofurantoin resistance, even in an additive way: E58D; I117T; K141E; L157F; A172S; G187D; A188V. The following variants, however, were unable to complement to nitrofurantoin susceptibility, and had reduced NfsA activity in order of increasing effect: R203C (14% of wild type); H11Y (7%); W212R (3%); A112E (<1%); A112T (zero activity); A122T (zero activity). These final two variants were found alongside A34V, and whilst we did not have the A34V variant alone in our collection so did not test it, the fact that A-V is a highly conservative change and that A112E alone and A34V, A112T had essentially the same effect on NfsA activity (i.e. activity was almost zero) it was considered unlikely that A32V was necessary for this effect. But this possibility cannot be entirely excluded.

For NfsB, the following amino acid substitutions did not have a significant effect on enzyme function: G66D; M75I; V93A; A174E. The R121C variant could still complement nitrofurantoin to susceptibility (to 32 mg/L, compared with 8 mg/L for the fully functional variants) but NfsB activity was reduced to 16% of wild-type levels. It is likely that over-expression of this partially functional *nfsB* because of its presence on a high copy number vector, pK18, explained the borderline nitrofurantoin susceptible MIC seen, and that in wild-type copy number this allele is unlikely to be sufficiently active to complement to nitrofurantoin susceptibility. Hence this variant was considered to be of reduced function. In order of increasing effect the following additional reduced function variants could not complement to nitrofurantoin susceptibility: Q142H (MIC 64 mg/L; 10% of wild-type activity); F84S (MIC 64 mg/L; 9%); P163H (MIC 128 mg/L; <1%); W46R (MIC 128 mg/L; zero activity); K57E (MIC 128 mg/L; zero activity) and V191G (MIC 128 mg/L; zero activity). One variant carried E137Q, K165E, L172F and had zero activity. It is therefore not possible to determine which, if not all three, of these variations causes this loss of enzyme activity.

### Nitrofurantoin “pre-resistance” among urinary E. coli from primary care

Factoring in the results of the above in vitro analysis of recombinant clones allowed us to identify 2 additional nitrofurantoin susceptible urinary isolates with loss of NfsA, making 6/166 (3/6%) of isolates in total (**Table 2**). No additional NfsB loss was identified, but with the 3/166 (1.8%) isolates with NfsB loss (**Table 3**), in total 9/166 (5.4%) of these nitrofurantoin susceptible isolates carried a mutation in *nfsA* or *nfsB* associated with nitrofurantoin resistance. We consider these as “pre-resistant” isolates, since they are more likely to evolve to a fully resistant phenotype than is a currently functionally wild-type isolate. It is important to remember, however, that the nitrofurantoin susceptible isolates sequenced here are resistant to one or more of cefpodoxime, amoxicillin/clavulanate or ciprofloxacin. Accordingly, they are not representative of the overall primary care urinary *E. coli* population. To illustrate this point, in 2020, the proportion of all primary care urinary *E. coli* resistant to nitrofurantoin was 1.34% based on 17581 *E. coli* positive primary care urine samples sent to the Severn Pathology diagnostic laboratory. Of 176/200 sequenced urinary isolates reported here that are resistant to one or more of cefpodoxime, amoxicillin/clavulanate or ciprofloxacin, 10/176 (5.7%) are nitrofurantoin resistant (**Table 1**). Nonetheless, when factoring in the data concerning single loss of function variants (**Table S1**, **Table 2**, **Table 3**), 19/176 (10.8%) of isolates resistant to one or more of cefpodoxime, amoxicillin/clavulanate or ciprofloxacin, are either nitrofurantoin resistant or pre-resistant, as defined above.

### NfsA/B downregulation is a common cause of nitrofurantoin resistance in urinary E. coli carrying genes encoding functionally wild-type enzymes

Integration of the results of our functional analysis confirmed a double NfsA/B loss of function in 25/34 (73.5%) nitrofurantoin resistant isolates (Table 1). However, 6 carried loss of function in *nfsA* but *nfsB* alleles encoding variants that should be functional. Furthermore, 3 carried functional alleles in both *nfsA* and *nfsB*. We hypothesised that these additional isolates would have downregulation of NfsA and/or NfsB production. To test this, we first assayed enzyme activity directly from cell extracts made from the isolates themselves (**Table 4**). These assays were calibrated against the standard susceptibility testing control *E. coli* strain ATCC25922 and mutants derived from this having insertional inactivation of either *nfsA* or *nfsB*.

**Table 4:** NfsA/B enzyme activity in nitrofurantoin resistant urinary *E. coli* from primary care carrying at least one functionally wild-type NfsA/B variant. Green amino acid substitutions represent a wild-type like function; orange, function not individually tested; red, loss of function; ND, not determined; **, statistically significant difference from wild-type ATCC25922, if not zero (*p*-value < 0.05).

| Isolate | NfsA Variant | NfsA+B<br>activity | Calculated |  | NfsA<br>Abundance | NfsB Variant | NfsB Activity |
| --- | --- | --- | --- | --- | --- | --- | --- |
|  |  |  | NfsA<br>activity |  |  |  |  |
| ATCC25922 | I117T, K141E, G187D, A188V | 6.03 +/- 0.25 | 2.25 |  | 0.021 +/- 0.001 | G66D, V93A, A174E | 3.78 +/- 0.30 |
| ATCC25922 | ΔnfsA | 2.69 +/- 0.11** | 0 | | ND | G66D, V93A, A174E | 3.71 +/- 0.24 |
| ATCC25922 | I117T, K141E, G187D, A188V | 4.84 +/- 0.09 | 4.84 | | ND | ΔnfsB | 0 |
| 39677* | 187FS | 0 | 0 |  | 0 | G66D, V93A, A174E | 0 |
| 39995* | L157F | 0.40 +/- 0.23** | 0 |  | 0 | M75I, V93A | 1.13 +/- 0.04** |
| 40104* | A112E, I117T, K141E, G187D | 0.96 +/- 0.22** | 0 |  | 0 | G66D, V93A, A174E | 1.88 +/- 0.19** |
| 40107* | A34V, A112T, I117T, K141E, G187D | 0.72 +/- 0.14** | 0 |  | 0 | G66D, V93A, A174E | 1.13 +/- 0.19** |
| 40115* | deletion | 0.17 +/- 0.11** | 0 |  | 0 | G66D, V93A, A174E | 0.52 +/- 0.05** |
| 40116* | 204Stop | 0.13 +/- 0.13** | 0 |  | 0 | G66D, V93A, A174E | 0.37 +/- 0.19** |
| 53866* | WT | 0 | 0 |  | 0 | WT | 0 |
| 53898* | I117T, K141E, E178G, G187D | 0 | 0 |  | 0 | G66D, V93A, A174E | 0 |
| 55968 | 55FS | 1.28 +/- 0.48** | 0 |  | 0 | V93A | 1.46 +/- 0.13** |

As hypothesised, all nitrofurantoin resistant isolates with functionally wild-type NfsB variants had statistically significantly downregulated NfsB enzyme activity (**Table 4**). Three isolates had no detectable NfsB activity at all, and the remainder had reduction of approximately 50% to 90%. We do not have a specific assay for NfsA enzyme activity, but by factoring in the measured NfsB activity, it was possible to calculate the contribution of NfsA to our NfsA/B combined assay. All tested isolates had no detectable NfsA activity, and this was confirmed by LC-MS/MS proteomics. Four of the isolates with zero detectable NfsA had nonsense or frameshift mutations, with one having a deletion; so we expected no protein production. Interestingly, the two isolates with NfsA reduced function missense variants, confirmed incapable of complementing to nitrofurantoin susceptibility, also had no detectable NfsA protein production. This apparent downregulation could be due to co-regulation alongside NfsB production; alternatively, because the mutant protein is unstable and liable to degradation. Nonetheless, we confirmed that the nitrofurantoin resistant isolates with functionally wild-type NfsA variants all had no detectable NfsA protein or enzyme activity. It is apparent therefore, that 9/34 (26%) of the nitrofurantoin resistant urinary *E. coli* sequenced here, with functionally wild type NfsB or NfsA/B, have genetic changes that have downregulated NfsA/B activity in other ways

### NfsA/B downregulation is, in most cases, due to unknown regulatory mutations

Analysis of the whole genome sequences of the 200 urinary isolates studied above identified that *nfsA* is the first gene in a predicted three gene operon. *In silico* promoter analysis identified the most likely promoter (ATGATTn17TATTCT within control isolate ATCC25922; 7/12 matches to the consensus *E. coli* δ^70^ promoter) starting 106 nt upstream of the *nfsA* start codon (**Fig. 2**). Of 3 nitrofurantoin resistant isolates carrying functional *nfsA* alleles, but with no detectable NfsA enzyme activity, isolate 39995 has an IS*1* insertion 70 nt upstream of the *nfsA* start codon, which removed the *nfsA* promoter, explaining why there is no NfsA activity. In isolate 53898, there is a mutation in the promoter, weakening the −10 (TATTCT becomes GATTCT) and in isolate 53866, a mutation weakening the −35 (ATGATT becomes ATGCTT). Notably, however, these two weaker promoter variants could be found among many of the 176 nitrofurantoin susceptible isolates sequenced for this study, so it is unlikely that either has a significant effect, and it is further evidence of random genetic drift at this locus.

**Figure 2.**
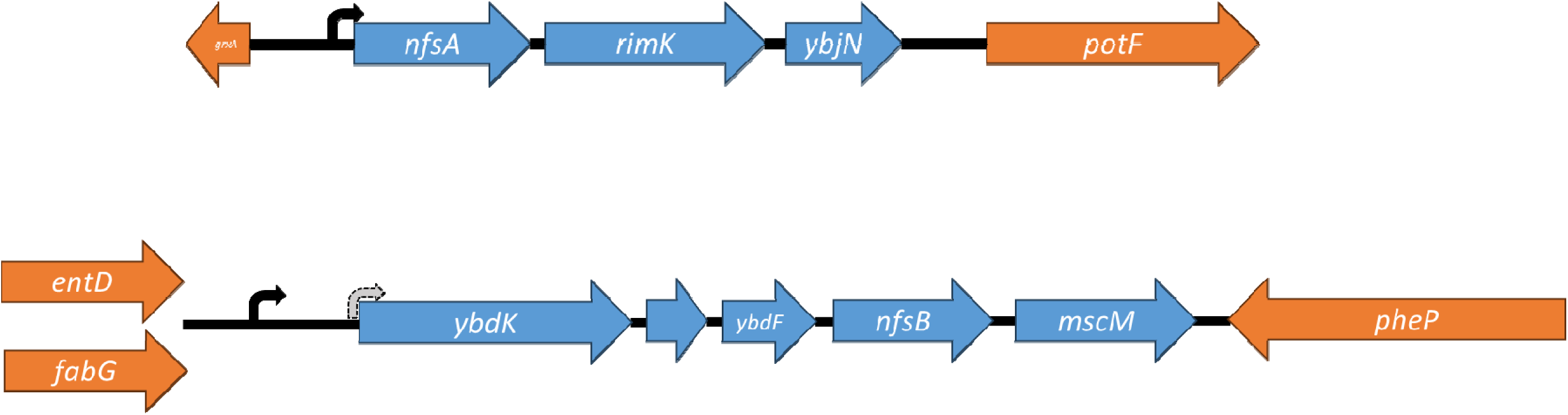
Schematic representation of the *nfsA* (top) and *nfsB* (bottom) loci found in 200 urinary *E. coli* isolates sequenced in this project. Promoter sequences were defined computationally and are noted as curved arrows. Where two possible promoters are noted for *nfsB*, the most likely based on match to the consensus promoter sequence is noted in black and the less likely match is noted in grey. Genes that form part of the *nfsA* or *nfsB* containing operons are marked in blue; flanking genes in orange. For *nfsB*, there are two alternative upstream sequences, marked. All genes and intergenic sequences (black lines) are drawn to scale and promoter positions are marked using the same scale.

*nfsB* is the fourth gene in a five gene operon with the first gene in this operon being *ybdK* (**Fig. 2**). The most likely promoter (TTGCCTn17TACCAT; 8/12 matches to the consensus *E. coli* δ^70^ promoter in control isolate ATCC25922) starts 443 nt upstream of *ybdK*. Based on the sequence further upstream, the isolates could be clustered into one of two types. Those with an intact *entD* gene upstream, and those with *entD* having been inactivated by insertion of 10 genes following a genomic rearrangement. The gene closest to *ybdK* in this second cluster of isolates is annotated as *fabG* (**Fig. 2**) though notably there are multiple variants of this gene in the *E. coli* genome and so apart from being a putative oxidoreductase its function is unknown.

Using the 5’ proximal 10 amino acids from the frameshifted versus the full length EntD amino acid sequence to text search genome annotation files, we counted that the *entD* upstream sequence for *ybdK* was found in 19/32 nitrofurantoin resistant and 70/165 susceptible isolates where the entire locus had not been deleted. This was not a statistically significant bias (chi squared 3.11; p=0.08). Furthermore, this variation in the upstream sequence does not affect the *ybdK* operon promoter, and so is unlikely to have any effect on *nfsB* expression. Of the nine isolates with confirmed reduced NfsB enzyme activity, only one had an obvious genetic lesion in the vicinity of *nfsB*: isolate 53866 has a 17.5 kb IS*26* mediated transposon insertion between *ybdK* and *nfsB*. This will disrupt transcription of *nfsB* as part of the operon, and is highly likely to explain why this isolate produces no NfsB. Isolate 55968 has a copy of IS*421* inserted between *ybdK* and *entD*, but this does not affect the *ybdK* operon promoter sequence. It remains possible, though unlikely, that this insertion has some impact on *nfsB* expression.

Overall, therefore, reduced NfsA and NfsB activity seen in the nitrofurantoin resistant isolates where genes are functionally wild-type appears mainly to be caused by genetic changes outside of the *nfsB* or *nfsA* loci, which remain to be identified.

## Discussion

There have been a number of surveys of nitrofurantoin resistant *E. coli* from clinical samples using PCR or in some cases whole genome sequencing to identify NfsA/B variants and in all cases a wide variety of mutations and strong evidence of random genetic drift has been observed (2–6). However, there has been little analysis of NfsA/B sequencing in nitrofurantoin susceptible isolates (6), and little attempt to definitively identify functionally wild-type versus reduced function variants. Indeed, in studies attempting to predict nitrofurantoin susceptibility based on whole genome sequence, the striking diversity of genes encoding NfsA and NfsB among the *E. coli* population have revealed a significant lack of concordance (6). Our perspective is that we do not yet know enough about the biology of NfsA/B function and control of production to be able to predict nitrofurantoin susceptibility from whole genome sequence. Instead we show how assay of NfsA/B and specific NfsB enzyme activity is necessary to define resistance. Ultimately deploying these assays will allow us to bridge the gap between nitrofurantoin susceptibility/resistance phenotype and whole genome sequence. Combined with complementation cloning, we show how enzyme assay can define which NfsA/B variants are functionally wild-type and which are not. Upon direct application of these assays to crude cell extracts of clinical isolates, we show very clearly that isolates having downregulated NfsA and NfsB are relatively common, even if they have functionally wild-type variants of these enzymes.

This is important, because there is interest in predicting nitrofurantoin susceptibility among surveillance collections not routinely tested for nitrofurantoin susceptibility – e.g. bloodstream isolates. This is for several reasons, not least that bloodstream isolates are more often sequenced than urinary isolates, and so surveillance for the emergence of nitrofurantoin resistant clones, or pre-resistance (where only one of NfsA or B are lost; something we have identified among primary care urinary *E. coli* here) within regional populations might be possible based on secondary analysis of these available datasets. Second, there is interest in whether nitrofurantoin resistance or even pre-resistance contributes to an increased odds of urosepsis, and in this case the isolates that must be surveyed for nitrofurantoin resistance might best be bloodstream isolates from suspected urinary versus gastrointestinal portal of entry into the blood. However, bloodstream isolates are not routinely tested for nitrofurantoin susceptibility because nitrofurantoin has no utility as a systemic antimicrobial. Given the huge number of possible NfsA/B sequence variants that might be encountered among routine surveillance studies (6), we suggest that concurrent NfsA/B activity assay should accompany whole genome sequencing in all cases where nitrofurantoin resistance is being considered. This is because as we have clearly demonstrated, phenotypic prediction is more likely to yield the correct result, since we have only just scraped the surface of defining the myriad possible NfsA/B reduced function and downregulated variants. Moreover an ever increasing number of comparisons between whole genome sequence and NfsA/B enzyme activity will provide the raw information needed to iteratively improve our understanding of genotype to phenotype relationships for nitrofurantoin resistance, hence its prediction.

## Materials and Methods

### Isolate collection and susceptibility testing

*E. coli* isolates were collected through routine diagnostic microbiology. These were from mid-stream urine samples sent in September and October 2020 from >150 primary care practices serving a population of approximately 1.5 million people in the English regions of Bristol, Bath, North Somerset and South Gloucestershire. Following diagnostic standard operating procedures, urine samples were plated onto ChromAgar Orientation and discs were directly applied. Zone diameters were interpreted by reference to EUCAST breakpoints (12). Isolates were collected if they were resistant to one or more of nitrofurantoin, cefpodoxime, ciprofloxacin and amoxicillin/clavulanate. Isolates were deduplicated by patient. All (n=34) nitrofurantoin resistant isolates collected during this time period plus the first sequential 166 nitrofurantoin susceptible isolates were collected. Nitrofurantoin MIC was defined using broth dilution methodology by following the standard EUCAST protocol. Resistance was defined using EUCAST breakpoints (12).

### Whole genome sequencing and bioinformatics

WGS was performed by MicrobesNG to achieve a minimum 30-fold coverage. Confluent growth for an isolate on TBX agar was streaked using a loop into a tube containing 10 mL of PBS to create a stock suspension. Density was adjusted by serial dilution and the optical density at 600 nm (OD_600_) was measured from the dilution point where OD_600_ was <1. The density of the stock solution was then adjusted based on this measured OD_600_ to be equivalent to an OD_600_ of 8 and 1 mL of the stock solution was transferred to an Eppendorf and the bacteria pelleted by centrifugation. The pellet was washed once in PBS and then resuspended in 500 µL of DNA/RNA Shield (Zymo Research) and shipped to MicrobesNG at ambient temperature.

Upon receipt, 5-40 µL of cell suspension were lysed with 120 µL of TE buffer containing lysozyme (0.1 mg/mL) and RNase A (0.1 mg/mL), by incubating for 25 min at 37°C. Proteinase K (0.1 mg/mL) and SDS (0.5% v/v) were then added followed by a further incubation for 5 min at 65°C. Genomic DNA was purified using an equal volume of solid-phase reversible immobilization beads and resuspended in 10 mM Tris-HCl, pH 8.0. DNA concentration was quantified with a Quant-iT dsDNA HS kit (ThermoFisher Scientific) assay in an Eppendorf AF2200 plate reader (Eppendorf UK Ltd, United Kingdom) and diluted as appropriate.

Genomic DNA libraries were prepared using the Nextera XT Library Prep Kit (Illumina, San Diego, USA) following the manufacturer’s protocol with the following modifications: input DNA was increased two-fold and PCR elongation time was increased to 45 s. DNA quantification and library preparation were carried out on a Hamilton Microlab STAR automated liquid handling system (Hamilton Bonaduz AG, Switzerland). Libraries were sequenced on an lllumina NovaSeq 2500 (Illumina, San Diego, USA) using a 250 bp paired end protocol.

Reads were processed using Trimmomatic version 0.39 (13). De novo assembly was performed using SPAdes version 3.13 (14). Assemblies were annotated using Prokka version 1.14.5 (15) and the annotation of AMR determinants was performed using ABRicate version 1.0.1 (https://github.com/tseemann/abricate), to search Resfinder including PointFinder (16). Sequence typing was performed using MOAST version 12.0 (17). NfsA/B variants were defined using the Hound, a bespoke pipeline for sequence analysis that uses tblastn to identify amino acid sequence variants relative to *E. coli* K-12 MG1655 as reference based on a de novo assembly of the genome (18). Putative promoter sequences were identified using BacPP (19).

### Phylogenetics

A core genome alignment and SNP tree were constructed using parSNP v1.5.3 with default parameters (20) to investigate phylogenetic relationships within this dataset, from the perspective of the core genome. The core genome SNP tree was overlaid with AMR data and visualised using iTOL version 5 (21).

### Insertional inactivation of nfsA and nfsB

*nfsA* and *nfsB* mutants were constructed by gene inactivation using the pKNOCK suicide plasmid (22) The DNA fragments were amplified with Phusion high-fidelity DNA polymerase (NEB, UK) from *E. coli* strain ATCC25922 genomic DNA. The DNA fragment of *nfsA* was amplified using primers *nfsA* forward (5’-TTT<u>ACTAGT</u>ATCGCTCCATTCGCCATTTC-3’) with *Spe*I restriction site, underlined, and *nfsA* reverse (5’-TTT<u>GTCGAC</u>CAGCGTCACCAGTTCTTCAC-3’) with *Sal*I site, underlined; and *nfsB* was amplified using primers *nfs*B forward (5’-TTT<u>ACTAGT</u>CAATACAGCCCATCCAGCAC-3’) with *Spe*I site, underlined, and *nfsB* reverse (5’-TTT<u>GTCGAC</u>TTTGCACAGAACACCACGAC-3’) with *Sal*I site, underlined. The PCR products were ligated to pKNOCK-GM at *Spe*I and *Sal*I sites yielded pKNOCK::GM-*nfsA* and pKNOCK::GM-*nfsB*, respectively. The recombinant plasmid was transferred by conjugation to *E. coli* ATCC 25922 carrying pK18 for counter selection (23). *nfsA* and *nfsB* mutants were selected on gentamicin (5 mg/L) and kanamycin (20 mg/L). Mutations were confirmed by PCR using primers *nfsA*-F (5’-ATA<u>GAATTC</u>ACGTCTTGCCCCACAGCTGATGA-3’) with *Eco*RI site, underlined, and BT543 (5’-TGACGCGTCCTCGGTAC-3’) or *nfsB*-F (5’-ATA<u>GAATTC</u>GAAATCTATAGCGCATTTTTCTC-3’) with *Eco*RI site, underlined, and BT543.

### Cloning nfsA and nfsB variants

*nfsA* and *nfsB* genes were amplified with Phusion high-fidelity DNA polymerase (NEB, UK) from various *E. coli* strain genomic DNA using primers *nfsA*-F with *Eco*RI site and *nfsA-*R (5’ ATA<u>GGATCC</u>ACATCGACGTGGCGGTTTTAGCGC-3’) with *Bam*HI site, underlined; or *nfsB*-F and *nfsB*-R (5’-ATA<u>GGATCC</u>GATGCCCGGCAAGAGAGAATTA-3’) with *Bam*HI site, underlined *nfsA* and *nfsB* variants were ligated into pK18 (24) at *Eco*RI and *Bam*HI restriction sites. pK18-*nfsA* and pK18-NfsB variants were transferred to isolate 39681 by electroporation and the recombinants were confirmed by kanamycin resistant and PCR with M13rev (5’-CAGGAAACAGCTATGAC-3’) and M13fwd (5’-GTAAAACGACGGCCAGT-3’) primers.

### Measurement of NfsA/B and NfsB enzyme activity

A 1.5 mL of overnight culture were pellet with 12,000 rpm for 1 min at room temperature. The pellet was resuspended with 75 µL of BugBuster® (Merck, Germany) and mixed at 300 rpm for 15 min at 20°C for cell lysis. Cell lysate was collected by centrifugation at 12,000 rpm for 20 min at 4°C. Ten µL of cell lysate was added to reaction mixture containing 10 mM Tris-HCl pH 7.0 with 50 mM NaCl, 100 µM Nitrofurantoin and 100 µM of NADPH (for NfsA/B activity) or 100µM of NADH (for NfsB activity) (11). The degradation of nitrofurantoin was measured at OD_400_ over 5 min for clinical isolates or over 1 min for *nfsA*/*nfsB* complementation; then enzyme specific activity were calculated as µmol min^-1^ ml^-1^ (ε_400_ = 7508 M^-1^ cm^-1^).

### Proteomic analysis

An overnight liquid culture was diluted to OD_600_ of 0.1 in 50 mL of fresh Cation Adjusted Mueller Hinton Broth and incubated at 37°C until an OD_600_ of 0.5 to 0.6. Samples were centrifuged at 4,000 rpm for 10 min at 4°C and the supernatants discarded. Cells were resuspended into 35 mL of 30 mM Tris-HCl, pH 8 and broken by sonication using a cycle of 1 s on, 1 s off for 3 min at amplitude of 63% using a Sonics Vibra-Cell VC-505TM. Non-lysed cells were pelleted by centrifugation at 8,000 rpm (Sorvall RC5B Plus using an SS-34 rotor) for 15 min at 4°C. The supernatant was used as a direct source of protein for analysis. LC-MS/MS was performed and data analysed as described previously (25) using 5 μg of protein for each run. Analysis was performed in triplicate, each from a separate batch of cells. Abundance for each test protein in a sample was normalized relative to the average abundance of ribosomal proteins in the same sample (25).

### Ethics Statement

This project is not part of a trial or wider clinical study requiring ethical review. All bacterial isolates came from routine diagnostic samples and no patient data was recorded or used.

## Data availability

Whole genome sequence data for the isolates reported in this study are deposited with the European Nucleotide Archive (https://www.ebi.ac.uk/ena/) under project accession number PRJEB72122.

## Acknowledgements

This work was funded by Medical Research Council grant MR/T005408/1, by grants MR/N013646/1, MR/S004769/1 and NE/N01961X/1 from the Antimicrobial Resistance Cross Council Initiative supported by the seven UK research councils and the National Institute for Health Research (NIHR), by NIHR Programme Grant for Applied Research NIHR204400, and by grant 82459 from the Welsh Government Rural Communities - Rural Development Programme 2014-2020 supported by the European Union and the Welsh Government. J.E.S and W.W.Y.L. both received a scholarship from the Medical Research Foundation National PhD Training Program in Antimicrobial Resistance Research (MRF-145-0004-TPG-AVISO). N.S. was supported by a postgraduate scholarship from the University of Bristol. We are grateful to laboratory staff at Severn Pathology, North Bristol NHS Trust, for collecting the isolates. WGS was performed by MicrobesNG, Birmingham, UK. The views expressed are those of the authors and not necessarily those of the NIHR or the Department of Health and Social Care.

