## Supplementary Information for "Improving Nitrofurantoin Resistance Prediction in *Escherichia coli* from Whole Genome Sequence by Integrating NfsA/B Enzyme Assays"

**Figure S1.** Details of 166 nitrofurantoin susceptible urinary *E. coli* isolates giving NfsA/B types and antibiotic susceptibility profiles.

| ISOLATE | NfsA Changes | | | | | | NfsA Type | NfsB Changes | | | | NfsB Type | nfsB Upstream | Cefalexin | Cefpodoxir | Ciprofloxac | Co-amoxicl | Gentamicir | Nitrofurant |
|---|---|---|---|---|---|---|---|---|---|---|---|---|---|---|---|---|---|---|---|
| 39970 | 117I -> T | 141K -> E | 187G -> D | | | | 1 | 66G -> D | 93V -> A | 174A -> E | | 1 | fabG | r | r | r | i | r | s |
| 39971 | 117I -> T | 141K -> E | 187G -> D | | | | 1 | 66G -> D | 93V -> A | 174A -> E | | 1 | entD | r | r | r | r | r | s |
| 39972 | 117I -> T | 141K -> E | 187G -> D | | | | 1 | 66G -> D | 93V -> A | 174A -> E | | 1 | fabG | r | r | r | r | r | s |
| 39973 | None | | | | | | WT | del entD to cysS | | | | | DEL | s | r | r | s | s | s |
| 39974 | 117I -> T | 141K -> E | 187G -> D | 228FS | | | | 66G -> D | 93V -> A | 174A -> E | | 1 | fabG | r | r | r | r | s | s |
| 39975 | 58E -> D | 117I -> T | 141K -> E | 172A -> S | 187G -> D | | 3 | 66G -> D | 93V -> A | 174A -> E | | 1 | entD | r | r | r | s | s | s |
| 39977 | 117I -> T | 141K -> E | 187G -> D | | | | 1 | 66G -> D | 93V -> A | 174A -> E | | 1 | fabG | r | r | r | s | i | s |
| 39982 | 117I -> T | 141K -> E | 187G -> D | | | | 1 | 66G -> D | 93V -> A | 174A -> E | | 1 | fabG | r | r | r | s | s | s |
| 39987 | 117I -> T | 141K -> E | 187G -> D | | | | 1 | 66G -> D | 93V -> A | 174A -> E | | 1 | fabG | r | r | r | r | s | s |
| 39988 | 117I -> T | 141K -> E | 187G -> D | | | | 1 | 66G -> D | 93V -> A | 174A -> E | | 1 | fabG | r | r | r | s | s | s |
| 39990 | 117I -> T | 141K -> E | 187G -> D | | | | 1 | 66G -> D | 93V -> A | 174A -> E | | 1 | fabG | r | r | r | s | s | s |
| 39991 | 117I -> T | 141K -> E | 187G -> D | | | | 1 | 66G -> D | 93V -> A | 174A -> E | | 1 | fabG | r | r | r | s | s | s |
| 39992 | 117I -> T | 141K -> E | 187G -> D | | | | 1 | 66G -> D | 93V -> A | 174A -> E | | 1 | fabG | r | r | r | s | s | s |
| 39996 | 117I -> T | 141K -> E | 187G -> D | | | | 1 | 66G -> D | 93V -> A | 174A -> E | | 1 | fabG | r | r | r | r | s | s |
| 39998 | 117I -> T | 141K -> E | 187G -> D | | | | 1 | 66G -> D | 93V -> A | 174A -> E | | 1 | fabG | r | r | r | s | s | s |
| 40003 | None | | | | | | WT | 66G -> D | 93V -> A | 174A -> E | | 1 | entD | r | r | r | s | s | s |
| 40004 | 117I -> T | 141K -> E | 187G -> D | | | | 1 | 66G -> D | 93V -> A | 174A -> E | | 1 | fabG | r | r | r | s | r | s |
| 40007 | 117I -> T | 141K -> E | 187G -> D | | | | 1 | 66G -> D | 93V -> A | 174A -> E | | 1 | fabG | r | r | r | r | r | s |
| 40008 | 117I -> T | 141K -> E | 187G -> D | | | | 1 | 66G -> D | 93V -> A | 174A -> E | | 1 | fabG | r | r | r | r | r | s |
| 40009 | 117I -> T | 141K -> E | 187G -> D | | | | 1 | 66G -> D | 93V -> A | 174A -> E | | 1 | fabG | r | r | r | r | s | s |
| 40011 | 117I -> T | 141K -> E | 187G -> D | | | | 1 | 66G -> D | 93V -> A | 174A -> E | | 1 | fabG | r | r | r | s | s | s |
| 40012 | 117I -> T | 141K -> E | 187G -> D | | | | 1 | 66G -> D | 93V -> A | 174A -> E | | 1 | fabG | r | r | r | s | s | s |
| 40014 | 117I -> T | 141K -> E | 187G -> D | | | | 1 | 66G -> D | 93V -> A | 174A -> E | | 1 | fabG | r | r | r | s | r | s |
| 40015 | 117I -> T | 141K -> E | 187G -> D | | | | 1 | 66G -> D | 93V -> A | 174A -> E | | 1 | fabG | r | r | r | r | r | s |
| 40016 | 117I -> T | 141K -> E | 187G -> D | | | | 1 | 66G -> D | 93V -> A | 174A -> E | | 1 | fabG | r | r | r | s | s | s |
| 40017 | 117I -> T | 141K -> E | 187G -> D | | | | 1 | 66G -> D | 93V -> A | 174A -> E | | 1 | fabG | r | r | r | r | s | s |
| 40018 | 117I -> T | 141K -> E | 187G -> D | | | | 1 | 66G -> D | 93V -> A | 174A -> E | | 1 | fabG | r | r | r | s | s | s |
| 40019 | 117I -> T | 141K -> E | 187G -> D | | | | 1 | 66G -> D | 93V -> A | 174A -> E | | 1 | entD | r | r | r | s | s | s |
| 40020 | 117I -> T | 141K -> E | 187G -> D | | | | 1 | 66G -> D | 93V -> A | 174A -> E | | 1 | fabG | r | r | r | s | s | s |
| 40022 | 117I -> T | 141K -> E | 187G -> D | | | | 1 | 66G -> D | 93V -> A | 174A -> E | | 1 | fabG | r | r | r | r | s | s |
| 40023 | 117I -> T | 141K -> E | 187G -> D | | | | 1 | 66G -> D | 93V -> A | 174A -> E | | 1 | fabG | r | r | r | s | s | s |
| 40024 | 117I -> T | 141K -> E | 187G -> D | | | | 1 | 66G -> D | 93V -> A | 174A -> E | | 1 | entD | r | r | r | s | s | s |
| 40025 | 117I -> T | 141K -> E | 187G -> D | | | | 1 | 66G -> D | 93V -> A | 174A -> E | | 1 | fabG | r | r | r | r | s | s |
| 40026 | 117I -> T | 141K -> E | 187G -> D | | | | 1 | 66G -> D | 93V -> A | 174A -> E | | 1 | fabG | r | r | r | s | s | s |
| 40027 | 117I -> T | 141K -> E | 187G -> D | | | | 1 | 66G -> D | 93V -> A | 174A -> E | | 1 | fabG | r | r | r | s | s | s |
| 40030 | 117I -> T | 141K -> E | 187G -> D | | | | 1 | 66G -> D | 93V -> A | 174A -> E | | 1 | fabG | r | r | r | s | s | s |
| 40031 | 117I -> T | 141K -> E | 187G -> D | | | | 1 | 66G -> D | 93V -> A | 174A -> E | | 1 | fabG | r | r | r | s | s | s |
| 40033 | 117I -> T | 141K -> E | 187G -> D | | | | 1 | 66G -> D | 93V -> A | 174A -> E | | 1 | fabG | r | r | r | r | r | s |
| 40034 | 117I -> T | 141K -> E | 187G -> D | | | | 1 | 66G -> D | 93V -> A | 174A -> E | | 1 | fabG | r | r | r | r | s | s |
| 40035 | 117I -> T | 141K -> E | 187G -> D | | | | 1 | 66G -> D | 93V -> A | 174A -> E | | 1 | fabG | s | r | r | s | s | s |
| 40037 | 117I -> T | 141K -> E | 187G -> D | | | | 1 | 66G -> D | 93V -> A | 174A -> E | | 1 | entD | r | r | r | r | s | s |
| 40039 | 58E -> D | 117I -> T | 141K -> E | 172A -> S | 187G -> D | | 3 | 66G -> D | 93V -> A | 174A -> E | | 1 | entD | r | r | r | r | s | s |
| 40043 | 117I -> T | 141K -> E | 187G -> D | | | | 1 | 66G -> D | 93V -> A | 174A -> E | | 1 | fabG | r | r | r | r | s | s |
| 40044 | 117I -> T | 141K -> E | 187G -> D | | | | 1 | 66G -> D | 93V -> A | 174A -> E | | 1 | fabG | r | r | r | r | r | s |
| 40045 | 117I -> T | 141K -> E | 187G -> D | | | | 1 | 66G -> D | 93V -> A | 174A -> E | | 1 | fabG | r | r | r | s | s | s |
| 40049 | 117I -> T | 141K -> E | 187G -> D | | | | 1 | 66G -> D | 93V -> A | 174A -> E | | 1 | fabG | r | r | r | r | s | s |
| 40117 | 112A -> E | 117I -> T | 141K -> E | 187G -> D | | | 6 | 66G -> D | 93V -> A | 174A -> E | | 1 | fabG | s | s | r | s | s | s |

| | | | | | | | | | | | | | | | | | | | |
|---|---|---|---|---|---|---|---|---|---|---|---|---|---|---|---|---|---|---|---|
| 53754 | 117I -> T | 141K -> E | 187G -> D | 188A -> V | | | 2 | 66G -> D | 93V -> A | 174A -> E | | 1 | entD | s | s | s | r | s | s |
| 53755 | 58E -> D | 117I -> T | 141K -> E | 172A -> S | 187G -> D | | 3 | 66G -> D | 93V -> A | 174A -> E | | 1 | entD | s | s | s | r | s | s |
| 53756 | 117I -> T | 141K -> E | 187G -> D | | | | 1 | 66G -> D | 93V -> A | 174A -> E | | 1 | fabG | s | s | s | r | s | s |
| 53757 | 117I -> T | 141K -> E | 187G -> D | | | | 1 | 66G -> D | 93V -> A | 174A -> E | | 1 | fabG | s | s | s | r | s | s |
| 53759 | 117I -> T | 141K -> E | 187G -> D | | | | 1 | 66G -> D | 93V -> A | 174A -> E | | 1 | entD | s | s | s | r | s | s |
| 53762 | 117I -> T | 141K -> E | 187G -> D | 188A -> V | | | 2 | 66G -> D | 93V -> A | 174A -> E | | 1 | entD | s | s | s | r | s | s |
| 53763 | None | | | | | | WT | 66G -> D | 93V -> A | 174A -> E | | 1 | entD | s | s | s | r | s | s |
| 53767 | 117I -> T | 141K -> E | 187G -> D | | | | 1 | 66G -> D | 93V -> A | 174A -> E | | 1 | fabG | s | s | s | r | s | s |
| 53768 | 117I -> T | 141K -> E | 187G -> D | 188A -> V | | | 2 | 66G -> D | 93V -> A | 174A -> E | | 1 | entD | s | s | s | r | s | s |
| 53775 | None | | | | | | WT | 66G -> D | 93V -> A | 174A -> E | | 1 | entD | r | s | s | r | s | s |
| 53779 | None | | | | | | WT | 66G -> D | 93V -> A | 174A -> E | | 1 | entD | s | s | s | r | s | s |
| 53781 | 117I -> T | 141K -> E | 187G -> D | | | | 1 | 66G -> D | 93V -> A | 174A -> E | | 1 | entD | s | s | s | r | s | s |
| 53784 | 117I -> T | 141K -> E | 187G -> D | 188A -> V | | | 2 | 66G -> D | 93V -> A | 174A -> E | | 1 | entD | s | s | s | r | s | s |
| 53785 | 117I -> T | 141K -> E | 187G -> D | | | | 1 | 66G -> D | 93V -> A | 174A -> E | | 1 | fabG | s | s | s | r | s | s |
| 53786 | 58E -> D | 117I -> T | 141K -> E | 172A -> S | 187G -> D | | 3 | 66G -> D | 93V -> A | 174A -> E | | 1 | entD | s | s | s | r | s | s |
| 53793 | None | | | | | | WT | 66G -> D | 93V -> A | 174A -> E | | 1 | entD | s | s | s | r | s | s |
| 53794 | None | | | | | | WT | 66G -> D | 93V -> A | 174A -> E | | 1 | entD | s | s | s | r | s | s |
| 53795 | 117I -> T | 141K -> E | 187G -> D | | | | 1 | 66G -> D | 93V -> A | 174A -> E | | 1 | fabG | s | s | s | r | s | s |
| 53798 | None | | | | | | WT | 66G -> D | 93V -> A | 174A -> E | | 1 | entD | s | s | s | r | s | s |
| 53799 | 117I -> T | 141K -> E | 187G -> D | 188A -> V | | | 2 | 66G -> D | 93V -> A | 174A -> E | | 1 | entD | s | s | s | r | s | s |
| 53801 | 117I -> T | 141K -> E | 187G -> D | | | | 1 | 66G -> D | 93V -> A | 174A -> E | | 1 | fabG | s | s | s | r | s | s |
| 53803 | 117I -> T | 141K -> E | 187G -> D | | | | 1 | 66G -> D | 93V -> A | 174A -> E | | 1 | fabG | s | s | s | r | r | s |
| 53804 | 58E -> D | 117I -> T | 141K -> E | 187G -> D | | | 4 | 66G -> D | 93V -> A | 174A -> E | | 1 | entD | s | s | s | r | s | s |
| 53807 | 117I -> T | 141K -> E | 187G -> D | 188A -> V | | | 2 | 66G -> D | 93V -> A | 174A -> E | | 1 | entD | s | s | s | r | s | s |
| 53809 | 58E -> D | 117I -> T | 141K -> E | 172A -> S | 187G -> D | | 3 | 66G -> D | 93V -> A | 174A -> E | | 1 | entD | s | s | s | r | s | s |
| 53811 | 117I -> T | 141K -> E | 187G -> D | 188A -> V | | | 2 | 66G -> D | 93V -> A | 174A -> E | | 1 | entD | r | s | s | r | s | s |
| 53812 | 117I -> T | 141K -> E | 187G -> D | 188A -> V | | | 2 | 66G -> D | 93V -> A | 174A -> E | | 1 | entD | s | s | s | r | s | s |
| 53813 | 117I -> T | 141K -> E | 187G -> D | | | | 1 | 66G -> D | 93V -> A | 174A -> E | | 1 | fabG | s | s | s | r | s | s |
| 53814 | 117I -> T | 141K -> E | 187G -> D | | | | 1 | 66G -> D | 93V -> A | 174A -> E | | 1 | entD | s | s | s | r | s | s |
| 53818 | 117I -> T | 141K -> E | 187G -> D | 188A -> V | | | 2 | 66G -> D | 93V -> A | 174A -> E | | 1 | entD | s | s | s | r | s | s |
| 53820 | 117I -> T | 141K -> E | 187G -> D | | | | 1 | 66G -> D | 93V -> A | 174A -> E | | 1 | fabG | s | s | s | r | s | s |
| 53822 | 117I -> T | 141K -> E | 187G -> D | 188A -> V | | | 2 | 66G -> D | 93V -> A | 174A -> E | | 1 | entD | s | s | s | r | s | s |
| 53824 | 117I -> T | 141K -> E | 187G -> D | | | | 1 | 66G -> D | 93V -> A | 174A -> E | | 1 | entD | s | s | s | r | s | s |
| 53826 | 117I -> T | 141K -> E | 187G -> D | 188A -> V | | | 2 | 66G -> D | 93V -> A | 174A -> E | | 1 | entD | s | s | s | r | s | s |
| 53828 | 117I -> T | 141K -> E | 187G -> D | 188A -> V | | | 2 | 66G -> D | 93V -> A | 174A -> E | | 1 | entD | s | s | s | r | s | s |
| 53830 | 58E -> D | 117I -> T | 141K -> E | 172A -> S | 187G -> D | | 3 | 66G -> D | 93V -> A | 174A -> E | | 1 | entD | s | s | s | r | s | s |
| 53839 | 117I -> T | 141K -> E | 187G -> D | 188A -> V | | | 2 | 66G -> D | 93V -> A | 174A -> E | | 1 | entD | r | s | s | r | s | s |
| 53841 | 117I -> T | 141K -> E | 187G -> D | | | | 1 | 66G -> D | 93V -> A | 174A -> E | | 1 | fabG | s | s | s | r | s | s |
| 53842 | 117I -> T | 141K -> E | 187G -> D | 188A -> V | | | 2 | 66G -> D | 93V -> A | 174A -> E | | 1 | entD | s | s | s | r | s | s |
| 53843 | 117I -> T | 141K -> E | 187G -> D | 188A -> V | | | 2 | 66G -> D | 93V -> A | 174A -> E | | 1 | entD | s | s | s | r | s | s |
| 53844 | 117I -> T | 141K -> E | 187G -> D | | | | 1 | 66G -> D | 93V -> A | 174A -> E | | 1 | fabG | s | s | r | s | r | s |
| 53846 | 117I -> T | 141K -> E | 187G -> D | | | | 1 | 66G -> D | 93V -> A | 174A -> E | | 1 | fabG | s | s | r | i | r | s |
| 53850 | 58E -> D | 117I -> T | 141K -> E | 172A -> S | 187G -> D | | 3 | 66G -> D | 93V -> A | 174A -> E | | 1 | entD | s | s | r | s | i | s |
| 53853 | 117I -> T | 141K -> E | 187G -> D | | | | 1 | 75M -> I | 93V -> A | | | 2 | fabG | s | s | r | r | r | s |
| 53854 | 117I -> T | 141K -> E | 187G -> D | | | | 1 | 75M -> I | 93V -> A | | | 2 | fabG | s | s | r | s | s | s |
| 53856 | 117I -> T | 141K -> E | 187G -> D | | | | 1 | 75M -> I | 93V -> A | | | 2 | fabG | s | s | r | s | r | s |
| 53858 | del rimK to rcdA | | | | | | | 75M -> I | 93V -> A | | | 2 | fabG | s | s | r | s | s | s |

| | | | | | | | | | | | | | | | | | | | |
|---|---|---|---|---|---|---|---|---|---|---|---|---|---|---|---|---|---|---|---|
| 53859 | 117I -> T | 141K -> E | 187G -> D | | | | 1 | 75M -> I | 93V -> A | | | 2 | fabG | s | s | r | s | s | s |
| 53861 | 117I -> T | 141K -> E | 187G -> D | | | | 1 | 93V -> A | | | | 3 | fabG | s | s | r | s | s | s |
| 53863 | None | | | | | | WT | 93V -> A | | | | 3 | entD | s | s | r | s | r | s |
| 53868 | 117I -> T | 141K -> E | 187G -> D | | | | 1 | 93V -> A | | | | 3 | fabG | s | s | r | s | s | s |
| 53870 | 117I -> T | 141K -> E | 187G -> D | | | | 1 | 93V -> A | | | | 3 | fabG | s | s | r | r | r | s |
| 53871 | 117I -> T | 141K -> E | 187G -> D | | | | 1 | 93V -> A | | | | 3 | fabG | s | s | r | s | s | s |
| 53875 | 117I -> T | 141K -> E | 187G -> D | | | | 1 | 93V -> A | | | | 3 | fabG | s | s | r | s | s | s |
| 53876 | 58E -> D | 117I -> T | 141K -> E | 172A -> S | 187G -> D | | 3 | 93V -> A | | | | 3 | entD | s | s | r | s | r | s |
| 53878 | 117I -> T | 141K -> E | 187G -> D | | | | 1 | 66G -> D | 75M -> I | 93V -> A | | 4 | fabG | s | s | r | s | s | s |
| 53880 | 117I -> T | 141K -> E | 187G -> D | 202T -> P | | | 7 | 66G -> D | 93V -> A | | | 5 | fabG | s | s | r | s | s | s |
| 53882 | ins IS26 | | | | | | | None | | | | WT | fabG | s | s | r | s | s | s |
| 53883 | 117I -> T | 141K -> E | 187G -> D | | | | 1 | None | | | | WT | fabG | s | s | r | s | s | s |
| 53886 | 117I -> T | 141K -> E | 187G -> D | | | | 1 | 66G -> D | 93V -> A | 174A -> E | | 1 | fabG | s | s | r | s | s | s |
| 53887 | 117I -> T | 141K -> E | 187G -> D | | | | 1 | 66G -> D | 93V -> A | 174A -> E | | 1 | fabG | s | s | r | s | r | s |
| 53888 | 117I -> T | 141K -> E | 187G -> D | | | | 1 | 66G -> D | 93V -> A | 174A -> E | | 1 | fabG | s | s | r | s | r | s |
| 53890 | 117I -> T | 141K -> E | 187G -> D | | | | 1 | 66G -> D | 93V -> A | 174A -> E | | 1 | fabG | s | s | r | s | s | s |
| 53895 | 117I -> T | 141K -> E | 187G -> D | | | | 1 | 66G -> D | 93V -> A | 174A -> E | | 1 | fabG | s | s | r | s | s | s |
| 53900 | 58E -> D | 117I -> T | 141K -> E | 172A -> S | 187G -> D | | 3 | 66G -> D | 93V -> A | 174A -> E | | 1 | entD | s | s | r | s | s | s |
| 53904 | 117I -> T | 141K -> E | 187G -> D | | | | 1 | 66G -> D | 93V -> A | 174A -> E | | 1 | fabG | s | s | r | r | s | s |
| 53908 | 117I -> T | 141K -> E | 187G -> D | | | | 1 | 66G -> D | 93V -> A | 174A -> E | | 1 | fabG | s | s | r | r | s | s |
| 53909 | 117I -> T | 141K -> E | 187G -> D | | | | 1 | 66G -> D | 93V -> A | 174A -> E | | 1 | fabG | s | s | r | s | s | s |
| 53910 | 117I -> T | 141K -> E | 187G -> D | | | | 1 | 66G -> D | 93V -> A | 174A -> E | | 1 | fabG | s | s | r | s | s | s |
| 53912 | 117I -> T | 141K -> E | 187G -> D | | | | 1 | 66G -> D | 93V -> A | 174A -> E | | 1 | fabG | s | s | r | s | s | s |
| 53914 | 58E -> D | 117I -> T | 141K -> E | 172A -> S | 187G -> D | | 3 | 66G -> D | 93V -> A | 174A -> E | | 1 | entD | s | s | r | s | s | s |
| 53915 | 117I -> T | 141K -> E | 187G -> D | | | | 1 | 66G -> D | 93V -> A | 174A -> E | | 1 | fabG | s | s | r | r | s | s |
| 53917 | 117I -> T | 141K -> E | 187G -> D | | | | 1 | 66G -> D | 93V -> A | 174A -> E | | 1 | fabG | s | s | r | s | s | s |
| 53919 | 117I -> T | 141K -> E | 187G -> D | | | | 1 | 66G -> D | 93V -> A | 174A -> E | | 1 | fabG | s | s | r | s | r | s |
| 53920 | 117I -> T | 141K -> E | 187G -> D | | | | 1 | 75M -> I | 93V -> A | | | 2 | fabG | s | s | r | s | s | s |
| 53921 | 117I -> T | 141K -> E | 187G -> D | | | | 1 | 75M -> I | 93V -> A | | | 2 | fabG | s | s | r | s | s | s |
| 53923 | 117I -> T | 141K -> E | 187G -> D | | | | 1 | 93V -> A | | | | 3 | fabG | s | s | r | s | s | s |
| 53925 | 117I -> T | 141K -> E | 187G -> D | | | | 1 | 93V -> A | | | | 3 | fabG | s | s | r | s | s | s |
| 53926 | 117I -> T | 141K -> E | 187G -> D | | | | 1 | 93V -> A | | | | 3 | fabG | r | s | r | s | s | s |
| 53927 | 117I -> T | 141K -> E | 187G -> D | | | | 1 | 93V -> A | | | | 3 | fabG | s | s | r | s | s | s |
| 53928 | 117I -> T | 141K -> E | 187G -> D | | | | 1 | 66G -> D | 75M -> I | 93V -> A | | 4 | fabG | s | s | r | s | s | s |
| 53929 | 58E -> D | 117I -> T | 141K -> E | 172A -> S | 187G -> D | | 3 | None | | | | WT | entD | s | s | r | s | s | s |
| 53931 | 117I -> T | 141K -> E | 187G -> D | | | | 1 | 92FS | | | | | fabG | s | s | r | s | s | s |

| | | | | | | | | | | | | | | | | | | | |
|---|---|---|---|---|---|---|---|---|---|---|---|---|---|---|---|---|---|---|---|
| 55932 | 187G -> D | | | | | | 5 | 66G -> D | 93V -> A | 174A -> E | | 1 | entD | s | s | r | s | s | s |
| 55899 | 117I -> T | 141K -> E | 187G -> D | | | | 1 | 66G -> D | 93V -> A | 174A -> E | | 1 | fabG | r | r | s | r | i | s |
| 55901 | 117I -> T | 141K -> E | 187G -> D | 188A -> V | | | 2 | 66G -> D | 93V -> A | 174A -> E | | 1 | entD | r | r | s | i | s | s |
| 55902 | 117I -> T | 141K -> E | 187G -> D | | | | 1 | 66G -> D | 93V -> A | 174A -> E | | 1 | fabG | r | r | s | r | i | s |
| 55903 | 117I -> T | 141K -> E | 187G -> D | | | | 1 | 66G -> D | 93V -> A | 174A -> E | | 1 | fabG | r | r | s | i | r | s |
| 55904 | 117I -> T | 141K -> E | 187G -> D | 188A -> V | | | 2 | 66G -> D | 93V -> A | 174A -> E | | 1 | entD | r | r | s | r | s | s |
| 55906 | None | | | | | | WT | 66G -> D | 93V -> A | 174A -> E | | 1 | entD | r | r | s | s | s | s |
| 55911 | None | | | | | | WT | 66G -> D | 93V -> A | 174A -> E | | 1 | entD | r | r | s | s | r | s |
| 55914 | 117I -> T | 141K -> E | 187G -> D | | | | 1 | 75M -> I | 93V -> A | | | 2 | entD | r | r | s | r | s | s |
| 55918 | 117I -> T | 141K -> E | 187G -> D | | | | 1 | 75M -> I | 93V -> A | | | 2 | entD | r | r | s | r | s | s |
| 55920 | 117I -> T | 141K -> E | 187G -> D | | | | 1 | 93V -> A | | | | 3 | fabG | r | r | s | s | s | s |
| 55921 | 117I -> T | 141K -> E | 187G -> D | 188A -> V | | | 2 | 93V -> A | | | | 3 | entD | r | r | s | r | s | s |
| 55922 | 117I -> T | 141K -> E | 187G -> D | | | | 1 | 93V -> A | | | | 3 | fabG | r | r | s | i | r | s |
| 55923 | 117I -> T | 141K -> E | 187G -> D | | | | 1 | 66G -> D | 75M -> I | 93V -> A | | 4 | entD | r | r | s | s | s | s |
| 55924 | 117I -> T | 141K -> E | 187G -> D | | | | 1 | 66G -> D | 93V -> A | 174A -> E | | 1 | entD | r | r | s | r | s | s |
| 55925 | 58E -> D | 117I -> T | 141K -> E | 172A -> S | 187G -> D | | 3 | 66G -> D | 93V -> A | 174A -> E | | 1 | entD | r | r | s | s | s | s |
| 55929 | None | | | | | | WT | 66G -> D | 93V -> A | 174A -> E | | 1 | entD | r | r | s | s | s | s |
| 55930 | 117I -> T | 141K -> E | 187G -> D | | | | 1 | 66G -> D | 93V -> A | 174A -> E | | 1 | entD | r | r | s | r | s | s |
| 55931 | 117I -> T | 141K -> E | 187G -> D | 188A -> V | | | 2 | 66G -> D | 93V -> A | 174A -> E | | 1 | entD | r | r | s | r | s | s |
| 55936 | None | | | | | | WT | 66G -> D | 93V -> A | 174A -> E | | 1 | entD | r | r | s | r | s | s |
| 55938 | 117I -> T | 141K -> E | 187G -> D | 188A -> V | | | 2 | 66G -> D | 93V -> A | 174A -> E | | 1 | entD | r | r | s | r | s | s |
| 55940 | None | | | | | | WT | 66G -> D | 93V -> A | 174A -> E | | 1 | entD | r | r | s | s | s | s |
| 55949 | 117I -> T | 141K -> E | 187G -> D | 188A -> V | | | 2 | 66G -> D | 93V -> A | 174A -> E | | 1 | entD | r | r | s | r | s | s |
| 55950 | 117I -> T | 141K -> E | 187G -> D | | | | 1 | 66G -> D | 93V -> A | 174A -> E | | 1 | fabG | r | r | s | s | s | s |
| 55951 | 117I -> T | 141K -> E | 187G -> D | | | | 1 | 66G -> D | 93V -> A | 174A -> E | | 1 | fabG | r | r | s | s | s | s |
| 55953 | 117I -> T | 141K -> E | 187G -> D | 188A -> V | | | 2 | 93V -> A | | | | 3 | entD | r | r | s | r | s | s |
| 55954 | None | | | | | | WT | 66G -> D | 93V -> A | | | 5 | entD | r | r | s | r | s | s |
| 55958 | 117I -> T | 141K -> E | 187G -> D | | | | 1 | 66G -> D | 93V -> A | 174A -> E | 183Y -> * | | fabG | r | r | s | s | r | s |
| 55959 | 58E -> D | 117I -> T | 141K -> E | 172A -> S | 187G -> D | | 3 | 66G -> D | 93V -> A | 174A -> E | | 1 | entD | r | r | s | s | s | s |
| 55961 | 117I -> T | 141K -> * | | | | | | 66G -> D | 93V -> A | 174A -> E | | 1 | entD | r | r | s | s | s | s |
| 55962 | 117I -> T | 141K -> E | 187G -> D | | | | 1 | 66G -> D | 93V -> A | | | 5 | fabG | r | r | s | s | s | s |
| 55965 | 117I -> T | 141K -> E | 187G -> D | | | | 1 | 66G -> D | 93V -> A | 174A -> E | | 1 | entD | r | r | s | r | s | s |
| 55966 | 117I -> T | 141K -> E | 187G -> D | | | | 1 | 66G -> D | 93V -> A | 174A -> E | | 1 | entD | r | r | s | r | s | s |
| 55969 | 117I -> T | 141K -> E | 187G -> D | 188A -> V | | | 2 | 66G -> D | 93V -> A | 174A -> E | | 1 | entD | r | r | s | s | s | s |
| 55972 | 117I -> T | 141K -> E | 187G -> D | 188A -> V | | | 2 | 66G -> D | 93V -> A | 174A -> E | | 1 | entD | r | r | s | r | s | s |
| 55978 | 58E -> D | 117I -> T | 141K -> E | 172A -> S | 187G -> D | | 3 | 66G -> D | 93V -> A | 174A -> E | | 1 | entD | r | r | s | s | s | s |
